# Neuronal phenotype defined by transcriptome-wide bursting kinetics in pyramidal and fast-spiking cells

**DOI:** 10.1101/2024.05.16.594495

**Authors:** Vanda Tukacs, Eva Kristine Fladhus, Dániel Mittli, Magor L. Lőrincz, James Eberwine, Katalin Adrienna Kékesi, Gábor Juhász

## Abstract

Single-cell sequencing revealed the transcriptional heterogeneity of neurons and introduced transcriptome-defined neuron types and subtypes. Temporal recordings of gene transcription showed that mRNA synthesis is arranged into transcriptional bursts which can be characterized by kinetic parameters. Here, we selected two distinct, functionally homogenous, and electrophysiologically well-defined neuronal cell types – the pyramidal cells and fast-spiking interneurons – for transcriptional burst analysis. Hierarchical clustering based on electrophysiological parameters recovered these cell types. In contrast, clustering based on transcripts failed to differentiate these neuronal phenotypes. We applied mathematical models to analyze burst kinetics of fast-spiking and pyramidal cells to explore the transcriptional underpinning of their distinct physiological functions. We found that inferred burst kinetic parameters are distinct measures of gene expression from average mRNA counts. Based on our results, we suggest that burst kinetics can be considered as a component of transcriptome-defined neuronal phenotype besides mRNA counts.

## INTRODUCTION

The inherent heterogeneity observed in brain cells underscores the necessity of phenotyping neurons to identify homogeneous cell groups, thereby enhancing the reproducibility of experimental data. Classical, morphology- and immunohistochemistry-based cell phenotyping have historically yielded significant advancements in neuroscience^1,2^. Recently, the advent of single-cell sequencing has generated vast datasets, unveiling tens of thousands of transcripts in differing abundances within individual cells. The application of hierarchical clustering to non-zero mRNA copy number data has expanded our understanding of transcriptome-defined neuron types or subtypes^3,4^. Additionally, injecting neuronal mRNAs into pluripotent cells has demonstrated the development of cells toward the donor cell’s phenotype^5^, suggesting a role for the cytoplasmic transcriptome in shaping cellular phenotypes.

High-temporal resolution measurements of gene transcription in bacteria^6,7^, yeast^8,9^, and mammalian cells^10,11^ have introduced the two-state model, also known as telegraph model^12^. This model posits ON and OFF states of genes organized into transcriptional bursts, with burst frequency and burst size controlled by the genomic DNA/chromatin dynamics that are regulated in part by the availability of transcription factors and RNA polymerase II^13^. The model implies that the measured transcript copy number results from the interplay of transcriptional burst frequency, size, and mRNA degradation contributing to the stochastic mRNA availability that is integral to the cellular phenotype^14,15,16,17^. Currently, single-cell transcriptomics stands as the sole technique capable of offering a genome-wide perspective on transcriptional fluctuation between ON and OFF states for multiple genes in *ex vivo* brain tissues.

In this study, we endeavor to model transcriptome-wide burst kinetics for classically described, functionally homogeneous neuron types, drawing upon electrophysiological recordings of their response characteristics. Single-cell RNA (scRNA) datasets, distinct from bulk transcriptomics studies, present snapshots at the time of the cell harvesting, with a substantial number of zero expression values representing a mix of biological and non-biological zeros^18^. This inherent variability, compounded by transcriptional burst-induced noise, necessitates the application of burst kinetics models to characterize the bursting activity of different genes^19,20,21,22,23^. Here, we adopt the beta-Poisson model proposed by Vu et al.^22^ due to its suitability for datasets with at least 25 cells and its ability to capture the long-tailed behavior and bimodality of mRNA count distributions. The model parameters (alpha and beta) infer burst frequency and burst size, respectively, enabling comparative bursting kinetics analysis of genes. Notably, transcriptome-wide burst modeling has not been previously applied to neuronal cell types, making our study a pioneering effort in elucidating the distinctive role of neuronal phenotypes.

For our investigation, we selected two well-defined neuron types, pyramidal cells (Pyr) and fast-spiking interneurons (FS) from the prefrontal cortex (PFC), chosen for their classical identification based on electrophysiological properties^24^. Pyr cells exhibit varied firing patterns^25^ and single-cell transcriptomics^26,27^ suggesting potential subtypes, the underlying factors contributing to these differences remain unclear. Pyr cells, the principal cortical cells located in different cortical layers^2^ and FS interneurons, known for their inhibitory feedback to Pyr cells, exhibit distinct passive and active electrophysiological properties^28^. Leveraging single-cell transcriptomics, we aim to explore the molecular underpinnings of these differences in transcriptional burst kinetics, providing insights into the temporal regulation of newly synthesized proteins. Despite previous findings of substantial differences in cell surface molecules within the same dataset^29^, transcript copy numbers alone failed to sufficiently describe the transcriptomic basis of FS and Pyr cell phenotypes. However, their distinct electrophysiological parameters allowed successful recovery of FS and Pyr cell clusters, highlighting the functional homogeneity of the harvested cell types.

In anticipation of potential therapeutic applications, our focus on FS and Pyr cells in the PFC is grounded in their role in local and far-field gamma oscillatory coupling, crucial for various cognitive processes^30^. This study represents the first attempt at burst kinetics analysis for these functionally significant neuronal cell types, paving the way for a deeper understanding of their transcriptional dynamics and potential implications for future therapeutic interventions.

## MATERIAL AND METHODS

### Cell identification in acute brain slices

Cell identification was achieved through a combination of anatomical and physiological markers. Pyr cells were identified by the presence of a prominent apical dendrite and their pyramidal-shaped larger cell bodies. Specifically, cells were collected from the 3^rd^ and 5^th^ cortical layer layers to ensure the inclusion of pyramidal cells with distinct anatomical, electrophysiological, and computational properties. FS were identified based on the distinctive features of narrow spike width (<0.5 ms^31^) and the high firing rate in response to depolarization. The detailed methodological description of brain slice preparation, cell harvesting, and mRNA amplification and sequencing can be found in our earlier publication^29^. In total, n=84 cells were measured and sequenced, comprising n=59 Pyr cells and n=25 FS interneurons from the mouse PFC. This sampling strategy ensured a comprehensive representation of the identified cell types, allowing for a robust analysis of transcriptional burst kinetics in functionally distinct neuronal populations.

### Hierarchical clustering of FS and Pyr cells

To quantify parameters of electrophysiological signals a script developed for evaluation of patch-clamp data (Intracellular spike analysis, Cambridge Electronic Design, http://ced.co.uk/downloads/scriptsiganal) was used.

To test the homogeneity of the FS and Pyr cell clusters we performed a principal component analysis of the 22 parameters and it was followed by hierarchical clustering written in Python. We used Welsh statistics for cluster identification.

Hierarchical clustering based on electrophysiological parameters was described in our previous paper^29^, briefly: physiological data were organized in frames, where the First Frame was defined as the frame in which the cell produced action potentials throughout the entire depolarizing current pulse. The Last Frame was the depolarization step which induced the highest firing rate of the neuron. For clustering we used data from the First Frame. A principal component (PC) analysis was applied to electrophysiological parameters (depicted on Fig. 7) and we considered the first 3 PCs for the hierarchical agglomerative clustering.

Euclidean distance was considered in the 3D space (the 3 PCs being the coordinates) between the data points, and the Ward linkage criterion was applied. The method we used is called Ward’s minimum variance criterion and it minimizes the total within-cluster variance.

Hierarchical clustering of cells based on scRNA count data was performed in Python, using seaborn^32^, matplotlib^33^, pandas^34^, numpy^35^, and scipy^36^ packages. Standard deviation of mRNA counts was calculated using ‘std’ function, ranked in a descending order, and the top 100 genes were used for hierarchical clustering. Clustering was performed by ‘clustermap’ function of the ‘seaborn’ package with Ward method similarly to electrophysiology-based hierarchical clustering.

### Beta-Poisson modeling of burst kinetics

We applied the four-parameter beta-poisson model developed in R by Vu et al.^22^ to infer parameters of transcriptional burst kinetics. The modified Poisson model returns 4 parameters: alpha and beta which characterize the burst frequency and burst size, respectively, in addition to 2 scaling parameters; lambda1 and lambda2. Three functions from the BPSC GitHub repository were used (v0.99.2): ‘estimateBPMatrix’, ‘getBPMCnullmatrix’ and ‘getMCpval’^22^; repository can be found here: https://github.com/nghiavtr/BPSC. The first function takes a scRNA sequenced dataset as input, calculates burst parameters for these genes and runs a goodness-of-fit chi-squared (χ^2^) test. The second function takes the output from the first function as input, and generates Monte-Carlo (MC) null distributions for the list of beta-Poisson models. The final function returns the list of MCp-values generated by the second function. After the estimation of parameters, we filtered out genes that had MCp-values smaller than 0.05. All details of the applied model can be found in the original publication of Vu et al.^22^.

### Stochastic simulation of transcriptional bursts

Based on the inferred alpha and beta parameters of the beta-poisson model, we ran simulations of transcriptional bursts. We developed an R script to visualize burst frequency and burst size differences between cell types of selected genes on a pseudotime scale. The simulation was based on a MatLab implementation of the Gillespie algorithm to gene expression, where protein quantity was simulated in time^37^. We modified the implementation to simulate transcribed RNA levels in transcriptional ON and OFF states of genes. The rate of a gene switching ON (k_on_) was set to the normalized alpha parameter, and the rate of a gene switching OFF was calculated from ON and OFF state probabilities. The ON state probability was calculated based on the formula of *p_on_*(*t*) = 1 − *e*^−kon*t^, where k_on_ is the rate of the ON state and *t* is the maximal time. In the two-state model of transcription a gene is either in ON or OFF state, accordingly, OFF state probability was calculated as *p_off_*(*t*) = 1*-p_on_*(*t*). The rate of OFF state (k_off)_ was calculated based on the formula k_off_ = (-ln(1-*p_off_*(*t*)))/t. The RNA synthesis rate (k_s_) was the normalized beta multiplied by k_off_. The amplitude of transcription intensity depended on the beta parameter.

### Data analysis and statistics

Overall, our analytical approach involved a combination of established statistical methods and visualization tools, ensuring a robust examination of transcriptional burst kinetics in the selected neuronal cell types. The scRNA-seq dataset utilized in this study originated from the previously published work by Ravasz et al.^29^; GEO accession number: GSE135060). Our analyses involved the application of R programming language (version 4.3.0)^38^, and Python 3^39^. To preprocess the scRNA-seq dataset, we excluded genes expressed in less than 20% of cells in both cell types. Subsequently, essential metrics such as mean and median copy numbers, copy number density, and gene variance were computed, with zero values excluded using the ‘MatrixGenerics’ package (version 1.12.3)^40^. The statistical significance of these metrics was evaluated using the ‘stats’ package in R (version 4.3.0); statistical details of experiments can be found in the result section and figure legends; results were considered significant with *p*-values less than 0.05. Assumptions of statistical tests were validated. For visualization, we generated plots using the ‘ggplot2’ (version 3.4.4), ‘gplots’ (version 3.1.3), and ‘GGally’ (version 2.2.1) packages in R^41,42,43^. These packages collectively facilitated the creation of informative and visually appealing plots to illustrate our findings.

#### Correlation analysis

Correlations of copy number descriptive statistics measures and inferred burst kinetic parameters were calculated in R by ‘cor.test’ function of the ‘stats’ package with Spearman’s rank correlation method. Assumptions of Spearman’s correlations were tested; two variables were on a continuous scale and represent paired observations in all cases, monotonic relationship was confirmed on scatter plots.

#### Formulation of regression models

Regression models were performed in R with ‘lm’ function of ‘stats’ package. Multicolinearity of variables was tested and was under 0.8 in all cases, and constant variance of the residuals was confirmed on a scatter plot. As burst kinetic inference was found to have low accuracy on low expression genes, we decided to filter out genes with median copy number less than 2. We formulated regression models which included the effect of mean copy number (*x̄*), percentage of ON state cells (pON), median copy number (*x̆*) of genes, and their interactions on burst frequency (bf) in equation (1) and burst size (bs) in equation (2).

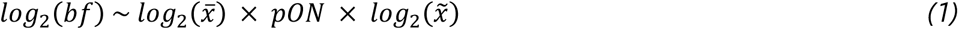

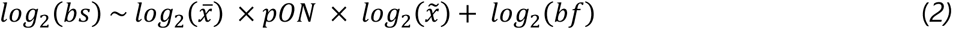

#### Differential burst kinetics

Next, we calculated alpha and beta parameter ratios of each gene by dividing parameters determined for FS interneurons and Pyr cells (FS/Pyr). The thresholds for differential burst frequency and burst size of genes were set as 3 times the standard deviation of the alpha and beta parameter ratios, respectively.

### Promoter motif analysis

For the analysis of promoter motifs, ensemble IDs of genes were used as input in the eukaryotic promoter database (EPD) selection tool^44,45^ (https://epd.expasy.org/epd/EPDnew_select.php), where promoters with TATA, GC, or CCAAT motifs were selected, for each gene, only the most representative promoter was selected. We ran the selection for 3 gene groups: all genes with burst kinetic parameters (All), genes that had larger burst frequency in FS interneurons (Alpha larger in FS), and genes that had larger burst size in Pyr cells (Beta larger in Pyr). Burst frequency and burst size values of genes with different promoter motifs were compared by Kruskal-Wallis test followed by pairwise comparisons using Wilcoxon rank sum test with continuity correction test in R with functions ‘kruskal.test’ and ‘pairwise.wilcox.test’ of ‘stats’ package. Next, we generated signal occurrence profiles of promoter motifs using the Oprof tool of EPD^45^. Relevant promoter regions were set as the following: TATA: from −80 position relative to transcription start site (TSS) to 20; CCAAT: from −200 to 20; GC: from −140 to 50. Occurrence profiles of promoter motifs of all genes with burst kinetic parameters were compared to frequency distributions of genes with alpha parameter larger in FS and genes with beta parameter larger in Pyr with two-sided Kolmogorov-Smirnoff test in R with ‘ks.test’ function of the ‘stats’ package. We integrated the relevant regions of the frequency plots - generated by Oprof - for each promoter motifs in OriginPro 9.0.0 to visualize promoter frequency differences between gene groups.

## RESULTS

### Hierarchical clustering of FS and Pyr cells based on single-cell transcriptome failed to recover cell-types

The identification of PFC cell types relied on electrophysiological parameters (see Supplementary fig. S1 A and B), discerned through principal component analysis. Hierarchical cluster analysis of the physiological data impeccably segregated FS and Pyr cells based on their distinctive electrophysiological signatures (Fig. 1A). In contrast, attempts to cluster FS and Pyr cells using the most variant 100 genes (Fig. 1B) resulted in the entwining of FS and Pyr cells within the final clusters. This observation implies transcriptome-level heterogeneity among FS and Pyr cells, emphasizing the inadequacy of mRNA copy numbers alone to distinguish physiological clusters of neurons. This discrepancy suggested that mRNA copy number data alone lacks the discriminatory power to delineate well-established, physiologically distinct neuronal phenotypes. Thus, our attention shifted towards exploring transcription dynamics, an essential parameter influencing mRNA availability for ribosomes and consequently impairing the translation of proteins.

**Figure 1.**
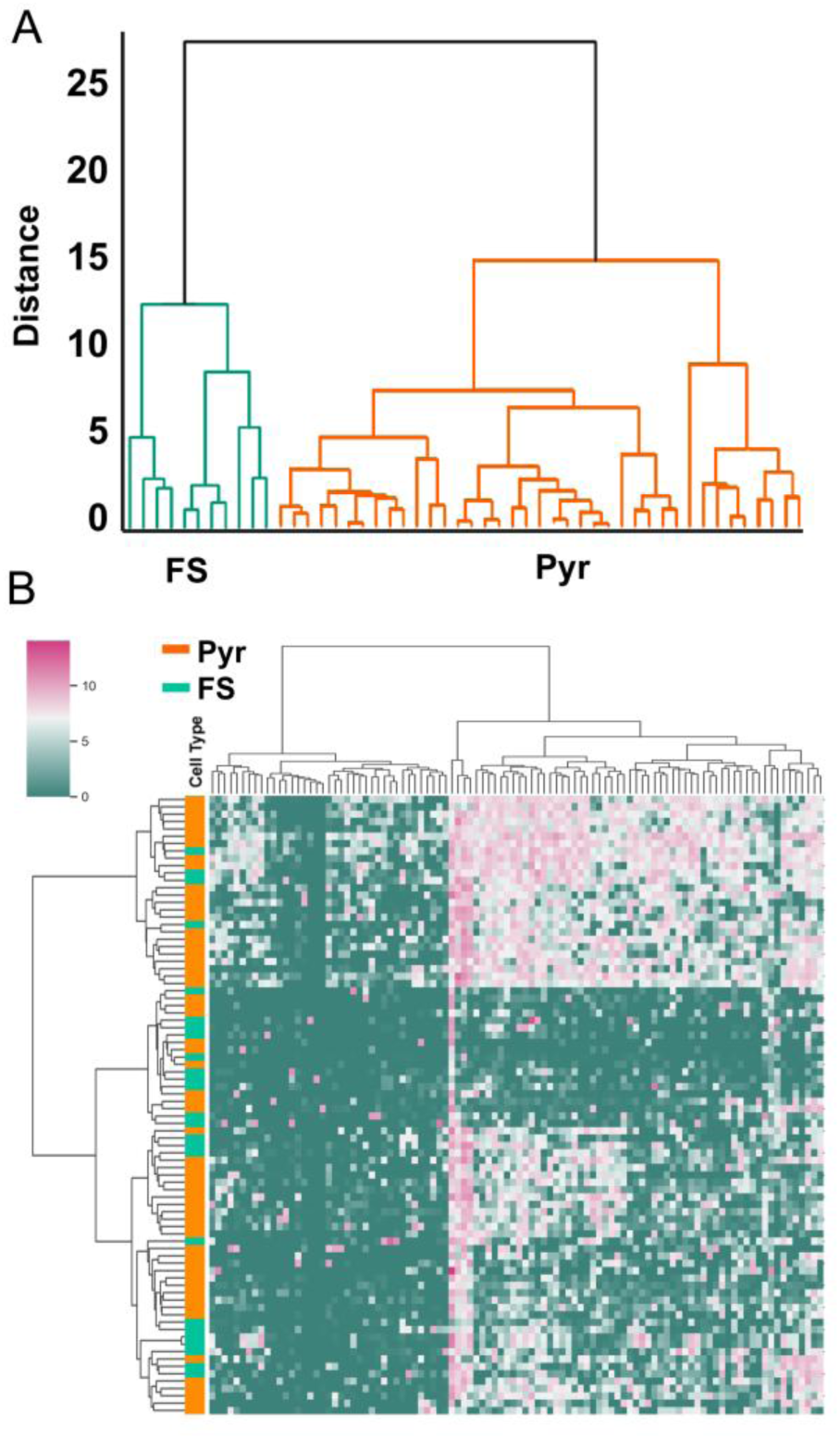
Hierarchical clustering of FS and Pyr cells based on single-cell transcriptome failed to recover cell-types. Hierarchical clustering based on electrophysiological data using four principal components successfully segregates Pyramidal (Pyr) and Fast-Spiking (FS) cell clusters (A), highlighting the distinct physiological signatures of these neuronal types. (B) In contrast, hierarchical clustering utilizing highly variant mRNA copy number of transcripts demonstrates the inability to recover distinct FS and Pyr cell clusters. The color scale represents the natural logarithm of copy numbers, illustrating the failure of transcriptome-based clustering to differentiate between FS and Pyr cells.

### Inferred burst kinetics parameters from beta-Poisson model suggest differential burst size between cell-types

The telegraph model of transcription postulates the existence of ON and OFF states of genes organized into transcriptional bursts with the measured copy number of a transcript being the outcome of transcriptional burst frequency, burst size, and RNA degradation process (Fig. 2A). Distinct transcriptional patterns in time emerge for genes with high or low burst frequencies and sizes (Fig. 2B).

**Figure 2.**
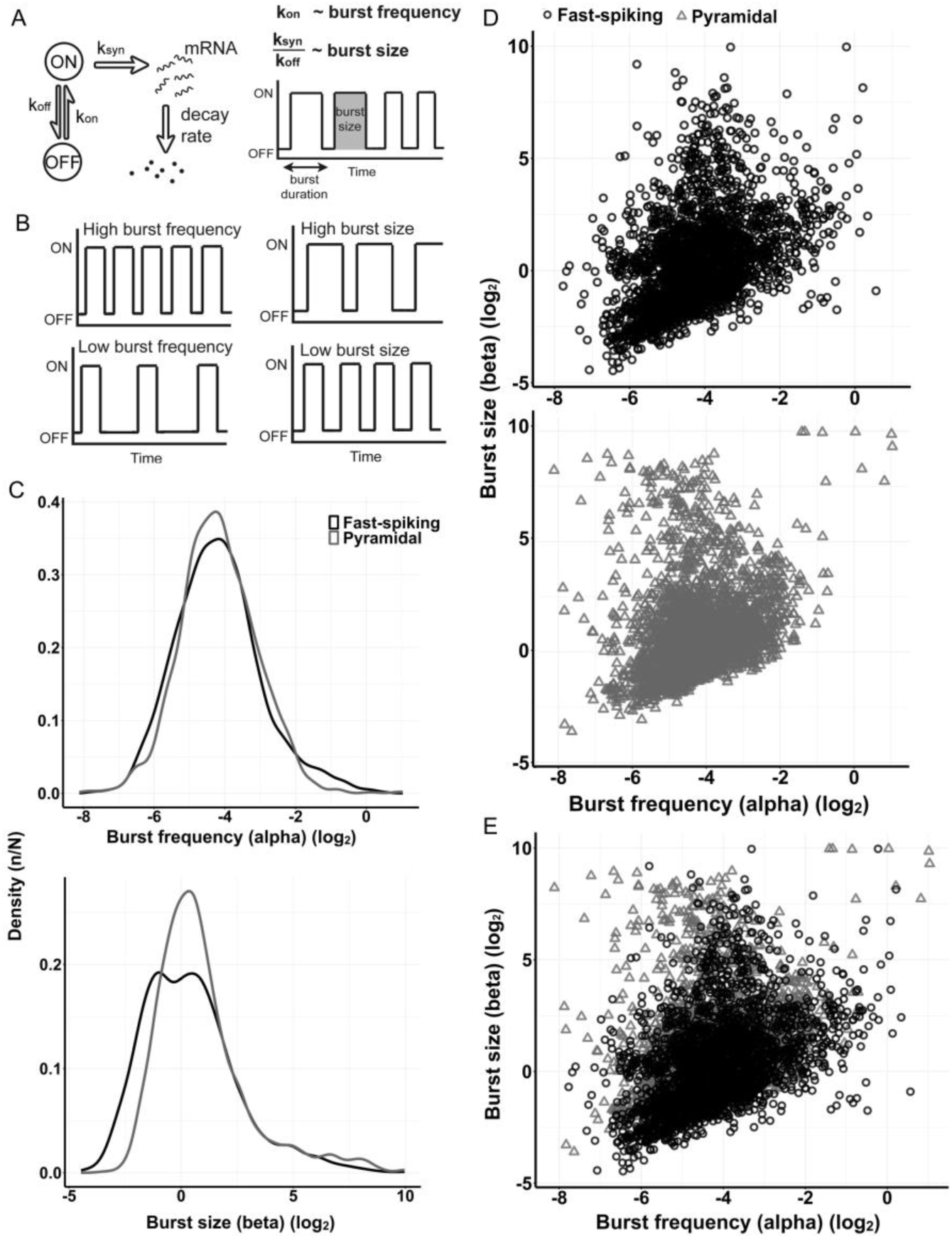
Inferred burst kinetics parameters from beta-Poisson model suggest differential burst size between cell-types. (A) Two-state model of transcription. A gene exhibits ON and OFF states with rates of k_on_ and k_off_, respectively. Transcription occurs at a rate of k_syn_ and the transcript subsequently decays. Burst frequency and burst size are characterized by k_on_ and k_syn_/k_off_, respectively. (B) Visual representations of burst frequency (left) and size (right) are shown. (C) Distribution of predicted alpha (upper) and beta (lower) parameters for genes in FS and Pyr cells. (D) Scatter plot of burst size in function of burst frequency in FS (upper) and Pyr (lower) cells. (E) Overlap of burst size in function of burst frequency for genes in FS and Pyr cells.

The beta-Poisson model of Vu et al.^22^ was originally tested on a breast cancer cell lines, colon cancer cell lines and simulated datasets but its application to neurons remained unexplored. The initial limit for the ON state number of a gene was set at 5% of the total number of cells, a criterion validated in previous datasets. However, when applied to our neuron datasets (59 Pyr cells and 25 FS interneurons), the 5% limit proved too conservative, resulting in very small parameter values for numerous transcripts.

Therefore, we filtered out genes expressed in less than 20% of the cells in either cell type. Next, we estimated beta-Poisson model parameters for all the selected genes using the ‘BPSC’ R package. Genes with Monte Carlo (MC) p-values less than 0.05 were discarded, yielding inferred burst kinetic parameters for 2630 genes in both cell types. The average alpha and beta values were 0.0795 ± 0.1031 and 9.99308 ± 50.79753 (mean ± sd), respectively, with beta exhibiting a larger standard deviation due to the substantial variation in copy numbers in the original data.

A comparison between FS and Pyr cells revealed wider alpha distribution with a longer tail in FS interneurons, while the alpha distribution in Pyr cells exhibited a higher and narrower peak. Although the alpha values did not exhibit a significant difference between cell types (two-sided Mann-Whitney test; n = 2630; *p* = 0.121; n indicates the number of genes) (Fig. 2C top), burst size (beta) displayed a bimodal distribution with double peaks. The density curve was wider in FS interneurons compared to Pyr cells, and the two exhibited a significant difference (two-sided Mann-Whitney test; n = 2630; *p* < 2.2e-16; n indicates the number of genes). The peak value of Pyr cell beta density plot was higher than FS beta (Fig. 2C bottom). In the inferred burst kinetics parameter space, genes in FS and Pyr cells predominantly occupied similar regions, forming a dense central patch in the plots with only a few genes showing separation (Fig. 2D and E).

### Average gene expression weakly correlates with inferred burst kinetics

Subsequent to the examination of the Spearman’s rank correlations for copy number features such as average, median, variance and the percentage of ON state cells, we observed robust correlations between average and variance copy numbers in both FS and Pyr cells (*ρ*_FS_ = 0.979; *ρ*_Pyr_ =0.981; n = 2630; n indicates the number of genes; Supplementary fig. S2A and B). Additionally, correlations of the percentage of ON state cells with other features were low in FS interneurons and moderate in Pyr cells, while correlations of median copy numbers were moderate to high in both FS and Pyr cells (Supplementary fig. S2A and B). Considering that burst frequency and burst size were inferred from mRNA counts, we anticipated a correlation between average mRNA copy numbers and inferred alpha and beta parameters (Fig. 3A). Surprisingly, correlations of average copy number with burst frequency (*ρ*_FS_ = −0.17; *ρ*_Pyr_ = 0.21; n = 2630) and burst size (*ρ*_FS_ = −0.068; *ρ*_Pyr_ = −0.25; n = 2630) were notably weak (n indicates the number of genes) (Fig. 3A, Supplementary fig. S2C and E). Although, a subtle trend emerged in the relation between burst frequency and the percentage of ON state cells (Supplementary fig. S2D), it was less apparent for burst size (Supplementary fig. S2F). To gain deeper insights, we formulated regression models to identify significant factors and interactions between burst kinetics and copy number distribution measures. Genes with median copy numbers less than 2 were filtered out due to the observed low accuracy of burst kinetic inference on low expression genes. Notably, for burst frequency (adj. *R*^2^ = 0.4705; *p*-value < 2.2e-16), average copy number, percentage of ON state cells, and their interactions emerged as significant coefficients. In the case of burst size (adj. *R*^2^ = 0.3788; *p*-value < 2.2e-16), burst frequency and median copy number were also significant factors, in addition to average copy number and percentage of ON state cells. Fitted burst frequency and burst size exhibited significant correlations with inferred burst frequency (*ρ*_FS_ = 0.651; *ρ*_Pyr_ = 0.75; n = 2630; *p*-value < 2.2e-16; Fig. 3B) and size (*ρ*_FS_ = 0.709; *ρ*_Pyr_ = 0.614; n = 2630; *p*-value < 2.2e-16; Fig. 3C), respectively (n indicates the number of genes). The high correlations between fitted and inferred kinetic parameters underscore the link between burst kinetics and not only average copy number but also median copy number, percentage of ON state cells, and – in the case of burst size - burst frequency of genes.

**Figure 3.**
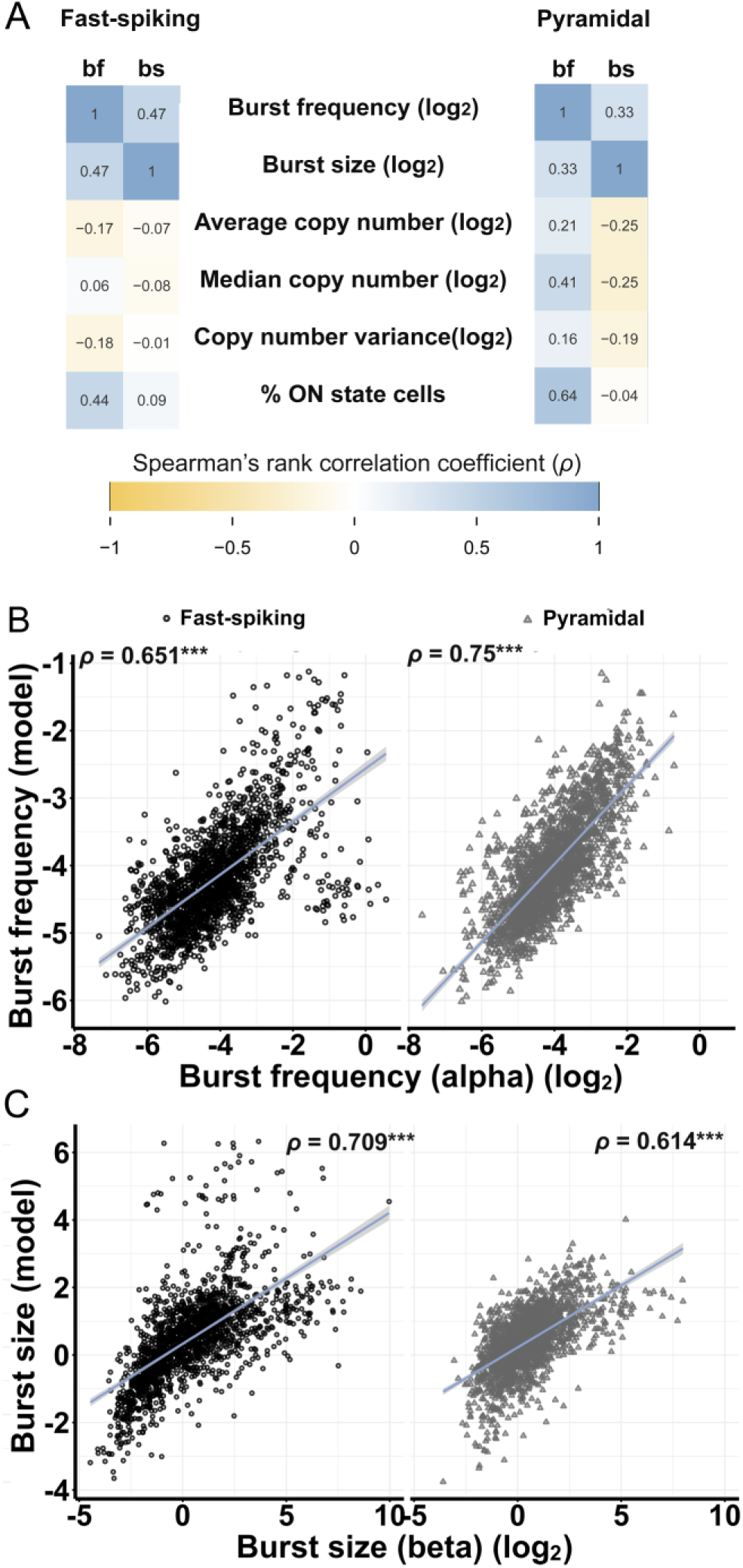
Average gene expression weakly correlates with inferred burst kinetics. (A) Spearman’s rank correlation between (log_2_ transformed) burst frequency (bf) and size (bs) with median, average copy number, copy number variance, and percentage of ON state cells in FS and Pyr cells. The color scale indicates the strength of the correlation. Correlations of inferred and fitted values (obtained from non-linear regression model) of (B) burst frequency and (C) size; Spearman’s rank correlation coefficients (*ρ*) are shown on the plot (***: *p*-value < 2.2e-16; n = 2630; n indicates the number of genes). (See also figure S2)

### Differential burst kinetics do not overlap with average copy number differences between FS and Pyr cells

To discern the transcriptional dynamics distinguishing FS and Pyr cells we evaluated burst kinetic parameters. The ratios of alpha (alpha_FS_/alpha_Pyr_) and beta (beta_FS_/beta_Pyr_) parameters were calculated to represent burst frequency and burst size differences between the two cell types. A threshold for differential burst kinetics was established as the threefold SD value of alpha and beta parameter ratios (Fig. 4A and B). Scatter plots of burst frequency against burst size highlighted genes exhibiting differential burst frequency and burst size in FS or Pyr cells (Fig. 4A and B). A total of 72 genes were identified with differential burst frequency of which 14 had larger burst frequency in Pyr cells and 58 in FS interneurons. Similarly, 66 genes displayed differential burst size, with 51 exhibiting larger burst size in Pyr cells and 15 in FS interneurons (Table 1, Supplemetary Table S3-S6).

**Figure 4.**
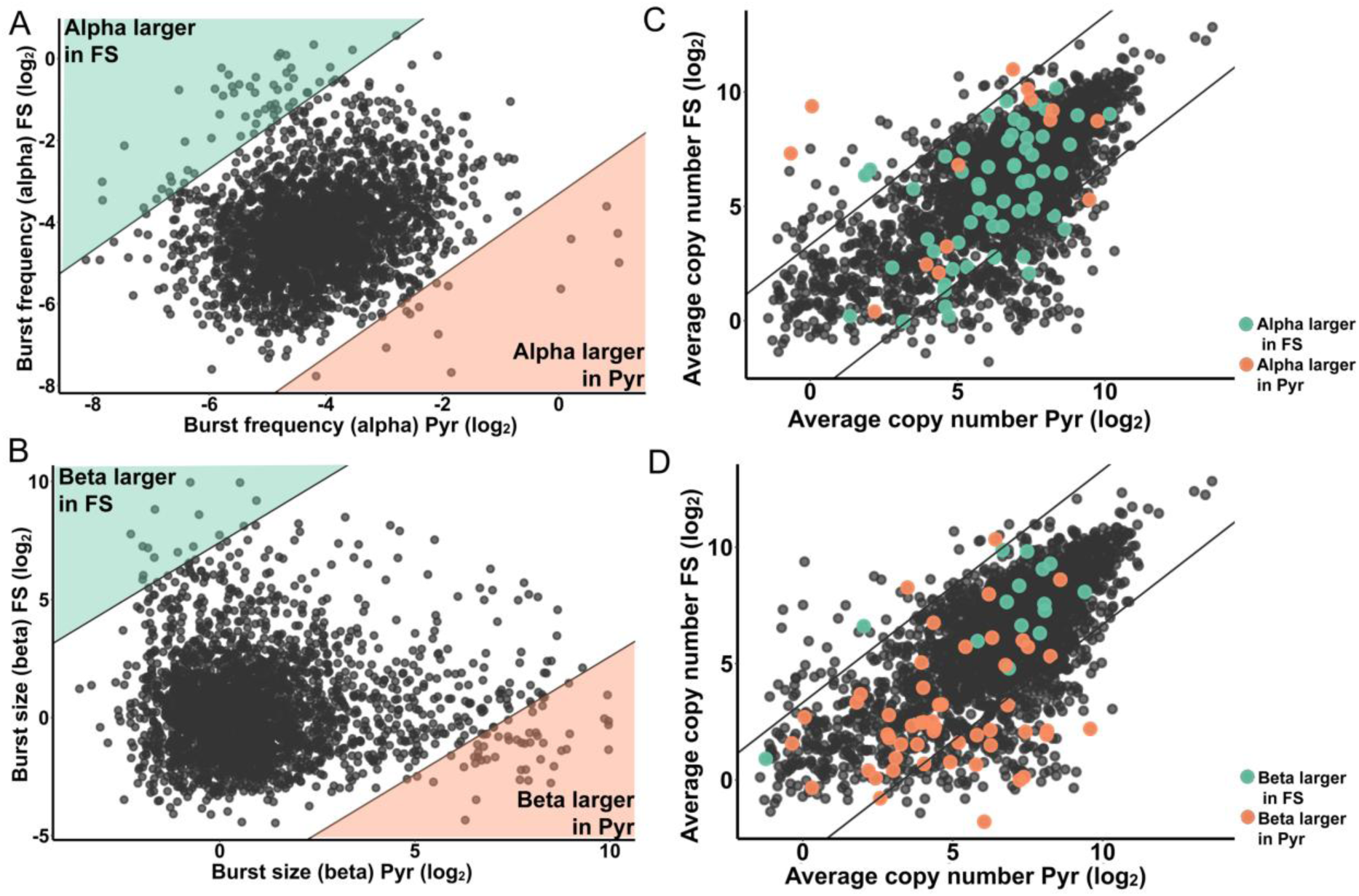
Differential burst kinetics parameters do not overlap with average copy number differences between FS and Pyr cells. (A) Burst frequency and (B) size plots of genes in FS and Pyr cells are presented; lines represent 3 times the SD of alpha and beta parameter ratios (FS/Pyr), respectively. Highlighted areas indicate genes with differential burst kinetic parameters. (C, D) Average copy number of genes in FS and Pyr cells are shown; lines correspond to 10 times differences in average copy numbers between cell types. Genes with differential (C) burst frequency and (D) size are highlighted (orange: parameter larger in Pyr cells; teal: parameter larger in FS).

**Table 1.**
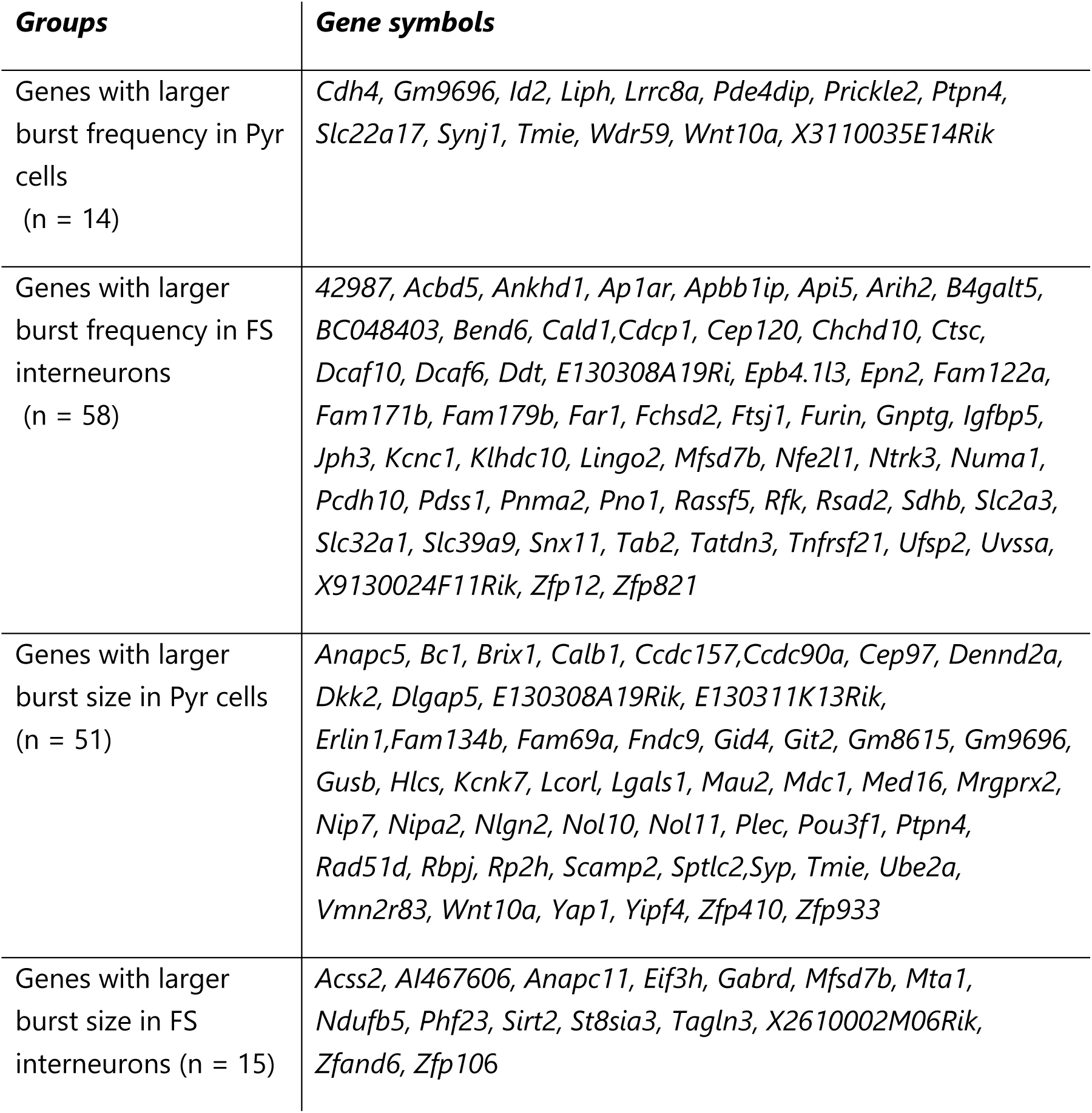
List of gene groups of differential inferred burst kinetics. (n indicates the number of genes) (See tables S3-S6.)

To elucidate the relationship between differential burst kinetics and average copy number variations, we highlighted the genes with differential burst frequency and burst size on the average copy number plot of FS and Pyr cells (Fig. 4C and D). This visual representation clearly illustrated the lack of overlap between differential alpha and beta values and differential copy numbers. Thus, our findings strongly suggest that the inferred burst kinetic parameters derived from the beta-Poisson model, based on scRNA sequencing, provide a unique insight into transcriptional dynamics.

We visualized the inferred bursting pattern in pseudotime of selected genes. Genes regulating physiological functions of FS and Pyr cells whose differential alpha or beta values according to the beta-Poisson model were chosen: solute carrier family 32 member 1 (*Slc32a1*), which is a vesicular amino acid transporter, gamma-aminobutyric acid receptor subunit delta (*Gabrd)*, voltage-gated potassium channel subfamily A member 1 (*Kcna1*), and potassium channel subfamily K member 7 (*Kcnk7*) (Fig. 5A). The transcriptional burst simulations of the selected genes depict their cell-type-specific bursting patterns. FS neurons exhibited a higher bursting frequency for the selected genes, while Pyr cells displayed larger burst sizes (Fig.5B).

**Figure 5.**
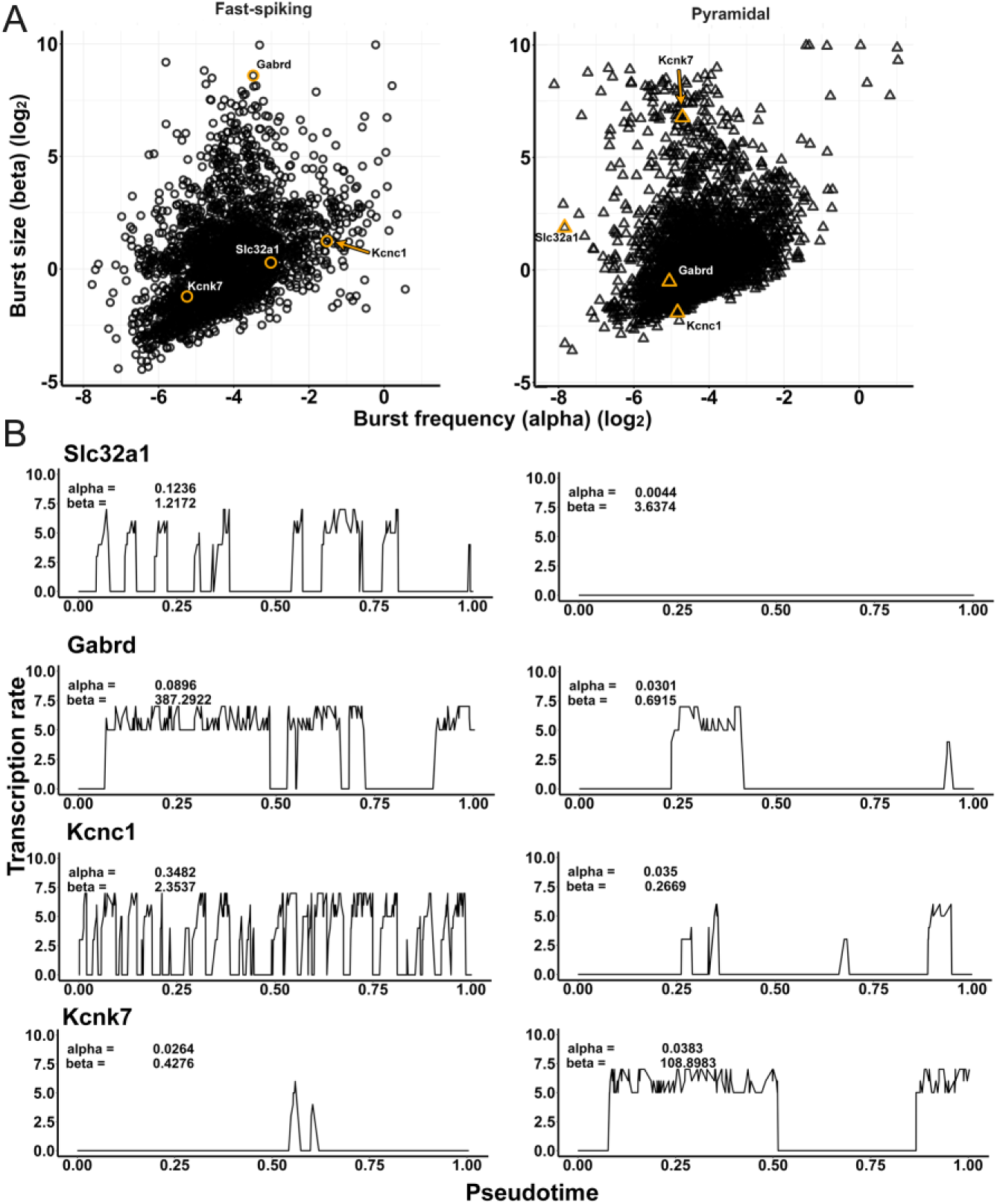
Stochastic simulations of transcriptional bursts of selected genes. (A) Selected physiologically important genes are highlighted in the scatter plot of burst size against burst frequency in FS (left) and Pyr (right) cells. (B) Stochastic simulations of transcriptional bursts on a pseudotime scale for the chosen genes are based on the inferred alpha and beta parameters (*Slc32a1*: solute carrier family 32 member 1 is a vesicular amino acid transporter; *Gabrd*: gamma-aminobutyric acid receptor subunit delta; *Kcna1*: voltage-gated potassium channel subfamily A member 1; *Kcnk7*: potassium channel subfamily K member).

### Inferred burst frequency is linked to promoter motif incidence

Transcription regulation involves intricate processes influenced by various factors, including chromatin modifications, DNA methylation, and transcription factor interactions with cis-regulatory elements as promoter regions (Fig. 6A). The assembly of the transcriptional machinery on the promoter, with RNA polymerase II initiating RNA synthesis at the transcription start site (TSS), is a crucial step. We sought to validate the impact of different promoter motifs on burst frequency or size.

**Figure 6.**
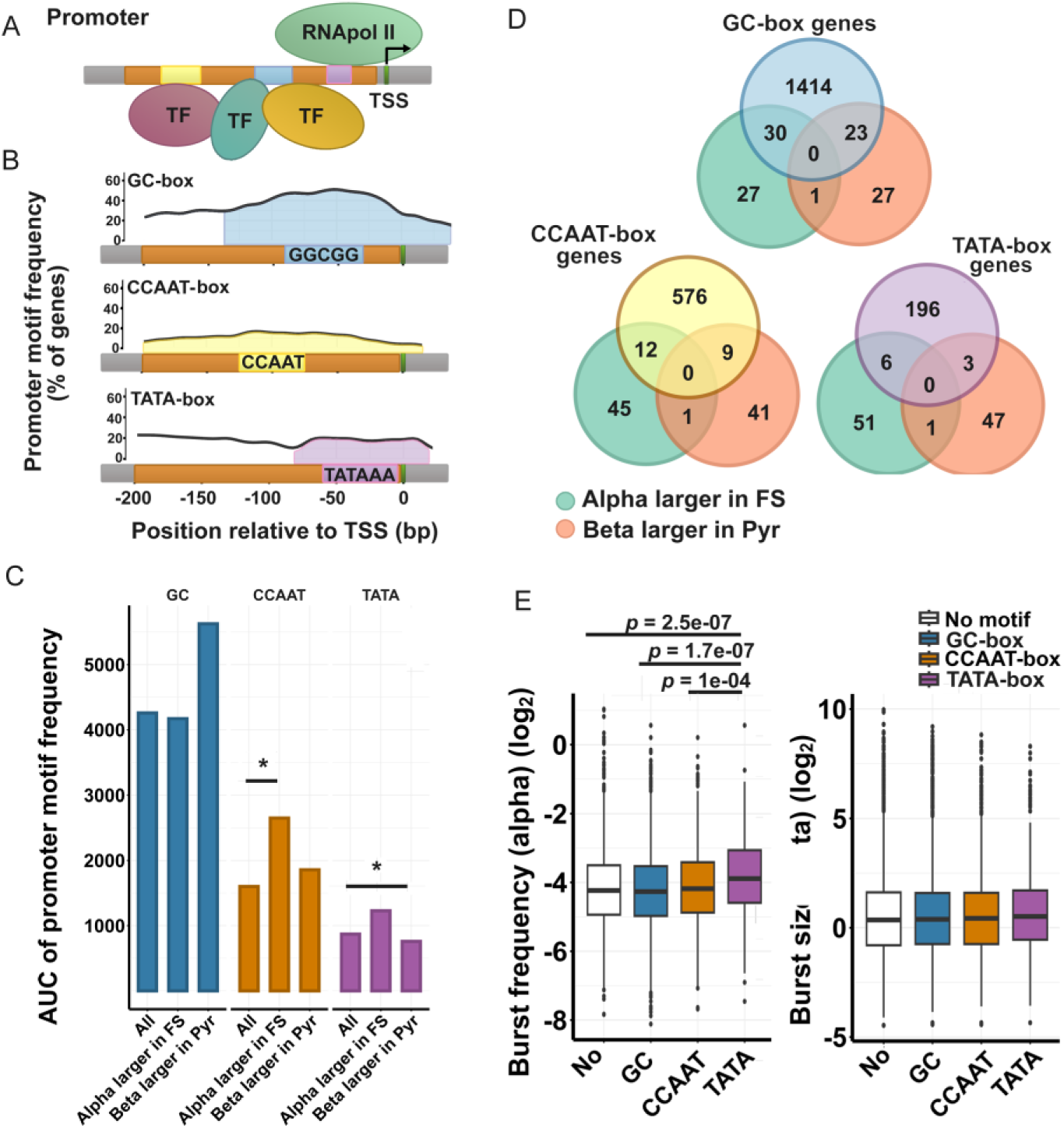
Inferred burst frequency is linked to promoter motif incidence. (A) Schematic illustration of the promoter region (TSS: transcription start site). (B) Analysis of GC-box, CCAAT-box, and TATA-box promoter motives in genes shows high frequency of GC-box. (C) Area under the curve (AUC) of promoter motif frequencies (*: *p* < 0.05 in Kolmogorov-Smirnoff test when relevant regions of promoter frequency distribution of genes were compared). (D) Overlap of genes with promoter motives and differential burst kinetic parameter. (E) Burst frequency and burst size distributions of genes with different promoter motives, significant *p*-values of pairwise Wilcoxon rank sum test are seen. (See also figure S3) (Boxes correspond to 1st and 3rd quartile, median is indicated by line; whiskers extend to minimum and maximal values but no longer than 1.5*inter-quartile range)

Using the EPD database, we identified genes with CCAAT- (n = 597), TATA- (n = 205), and GC-box (n = 1467) motifs, among the 2630 genes with inferred kinetic parameters (Fig. 6B) (n indicates the number of genes). The GC-box motif exhibited the highest frequency, while TATA- and CCAAT-box frequencies were comparatively lower. We also compared promoter motif frequencies between gene groups based on differential burst kinetics (Fig. 6C), comparing the relevant regions of promoter frequency distribution of genes with those of genes having alpha parameter larger in FS and beta parameter larger in Pyr (Supplementary fig. 3A). Notably, we excluded groups with low gene numbers from the analysis (genes with higher burst frequency in Pyr cells and genes with higher burst size in FS interneurons).

The analysis revealed that CCAAT-box motif frequency was higher in genes with higher burst frequency (alpha) in FS interneurons compared to all genes (*p*-value = 9.45e-03; Kolmogorov-Smirnoff test). Additionally, TATA-box motif frequency was significantly lower in genes with higher burst size (beta) in Pyr cells compared to all genes (*p*-value = 5e-04; Kolmogorov-Smirnoff test) (Fig. 6C). However, genes with differential burst kinetics and predicted promoter motifs in the EPD database constituted a small percentage—3.61%, 3.52%, and 4.39% of GC-, CCAAT-, and TATA-box genes, respectively (Fig 6D).

Further analysis suggested that burst frequency significantly differed between genes with differing promoter motifs analyzed by Kruskal-Wallis test (KW) (KW_bf_: Df = 3; *p*-value = 2.552e-07; KW_bs_: Df = 3; *p*-value = 0.5284). Pairwise Wilcoxon test revealed genes with TATA-box motif exhibited significantly higher inferred burst frequency compared to other motif groups (TATA-CCAAT: *p*-value = 1e-04; TATA-GC: *p*-value = 1.7e-07; TATA-No motif: *p*-value = 2.5e-07; Fig. 6E). A similar approach, considering gene groups with one or a combination of two promoter motifs, yielded consistent results (KW_bf_: Df = 5; *p*-value = 8.704e-05; KW_bs_: Df = 5; *p*-value = 0.7699; Supplementary fig. 3B). In conclusion, our findings suggest that genes with known TATA-box motifs, exhibit higher inferred burst frequencies.

## DISCUSSION

This study presents the first demonstration of transcriptome-wide burst kinetics as a component of FS and Pyr cell phenotypes. The analysis involved applying mathematical models to scRNA-seq data, allowing the estimation of burst size and burst frequency for a subset of genes in these cell types. The unique challenge in our investigation was the lack of a direct technique for transcriptome-wide analysis of burst kinetics in brain tissue, in contrast to the direct assessment possible for a few genes using single-molecule FISH (smFISH) technology^46,47,48^. Mathematical modeling based on scRNAseq data enabled transcriptome-wide burst size and burst frequency estimation, overcoming the absence of direct techniques^19,20,22,23^. Our application of the beta-Poisson model to neuronal datasets showed promise for capturing the long-tailed behavior and bimodality of mRNA count distribution, especially when dealing with a relatively small dataset.

### Considerations about modeling-based transcriptional burst analysis

In single-cell sequencing, the variability of mRNA copy numbers is higher compared to bulk tissue RNA sequencing studies, which may partly originate from transcriptional bursts. The main principle of modeling transcriptional burst kinetics on a cell group is that cell-to-cell variation of transcription - due to the heterogeneity of cellular transcriptome regulation - should be smaller than the variation derived from sampling the transcriptional bursts in different time points. Previously applied mathematical models are based on fitting a mixed-Poisson distribution on mRNA count distribution^19,20,22,23^. In most of the initial modeling studies, the authors validated the results by analogue measurement of gene transcription kinetics by smFISH technology, and revealed close correlation between modeling data with the results of real-time recordings^10,19^.

FS and Pyr cells are physiologically distinguishable in the functioning brain in a freely moving animal^24^. It is well known that Pyr cells can be clustered into subtypes on the basis of their responses to stimuli^25^ and based on single-cell sequencing^26,27^. FS interneurons could be selected properly by using immunohistochemically detectable markers as glutamate decarboxylase and parvalbumine (PV)^1^. However, in our analysis at transcription level, it was not possible to recover the FS and Pyr cell types on the basis of mRNA copy numbers by hierarchical clustering. In contrast, the hierarchical clustering on the basis of electrophysiological properties was successful, supporting that the selected cell types were functionally homogeneous.

### Are the burst size and burst frequency principal components of the cellular phenotype?

The main neuronal phenotypes as FS and Pyr cells can be classified by molecular and physiological markers^1,2^. Neuron typing was a great advance in neurosciences because it improved reproducibility of results and provided detailed insight into actual and potential function of said cells. Detection of neuron types is based on a snapshot so it is not clear whether the same cell stays in its suggested cell type in long term^49^. It is particularly true for great number of subtypes obtained on the basis of scRNA-seq data by clustering based on high number transcripts^26,27^.

Hierarchical clustering by scRNA-seq data groups cells with similar expression patterns into clusters. However, Dueck et al.^50^ revealed that within-cell-type variability is comparable to between-cell-type variability by single-cell sequencing analysis when analyzing scRNA-seq data of 5 distinct cell types from mice. Intriguingly, genes with high expression variability were enriched in functional categories that are relevant for their phenotypes in a cell-type-specific manner^50^. Considering this, highly variable genes may have more relevance in distinguishing cell types than genes with consistent expression within a cell type or genes with differential average copy numbers between cell types. Thus, identifying novel subtypes by performing hierarchical clustering or differential gene expression based on mRNA copy numbers in scRNA–seq studies might be misleading. Here, we chose a reversed approach, where we analyzed two functionally identified cell types of the mouse PFC and attempted to find genes that are transcribed differential burst kinetic parameters.

In our dataset, average copy numbers of genes did not correlate well with bursting kinetics in contrast to previous studies^19,20,51^. However, regression model of burst frequency, which included the percentage of ON state cells, average and median copy numbers highly correlated with inferred burst frequency. Similarly, regression model of burst size, which included burst frequency, the percentage of ON state cells, average, and median copy numbers highly correlated with inferred burst size. This indicates that inferred burst kinetics is linked to a group of measures characterizing copy number distributions. In addition, differences in burst frequency and burst size of particular genes in FS and Pyr cells did not overlap with the average copy number differences of the genes. It demonstrates that average copy number differences between the two neuron types are distinct from the transcriptional burst frequency and burst size differences. Further studies should address the biological relevance of transcriptional burst parameters as frequency and size and discover the correlation between protein availability and burst kinetics in neurons.

### Functional relevance of differential burst kinetics in FS and Pyr cells

Pyr cells integrate synaptic inputs collected by their reaching the superficial layers of the cerebral cortex, while FS interneurons perform an effective inhibitory feedback to the Pyr cells responsible for post-stimulus inhibition. This excitatory/inhibitory balance plays a crucial role in maintaining cortical circuits which are responsible for information processing and their perturbations drive several CNS disorders, such as seizure disorders, schizophrenia, autism spectrum disorder^52^, Alzheimer’s disease, Parkinson’s disease^53^, and amyotrophic lateral sclerosis^54^.

The GABAergic system - consisting of GABA, its transporters, andreceptors, and inhibitory synapses - is responsible for regulating Pyr cell activity via an inhibitory feedback feedback and feedforward mechanisms performed by GABAergic FS cells and other interneurons. Two genes of the GABAergic system were found to have differential burst kinetics, namely *Gabrd* and *Slc32a1*, which encode the delta subunit of GABA_A_ receptor and a vesicular GABA transporter, respectively. In FS interneurons, *Gabrd* and *Slc32a1* had higher burst size and burst frequency, respectively; their mutations were found in several types of seizure disorders^55,56^.

The sustained, high-frequency firing of FS neurons is enabled by *Kv3.1* (also known as *Kcnc1*) and *Kv3.2* voltage-gated potassium channels^57^; we found that *Kcnc1* has higher transcriptional burst frequency in FS interneurons. Heterozygous mutations of *KCNC1* in humans can cause seizure disorders, namely myoclonus epilepsy and ataxia (MEAK) due to potassium channel mutation^58^, and epileptic and developmental encephalopathy^59^. NT-3 growth factor receptor (*Ntrk3*) had also higher transcriptional burst frequency in FS interneurons; it is expressed by FS PV^+^ interneurons during synaptogenesis and its downregulation led to a decrease in the density of excitatory synapses received by PV^+^ interneurons^60^, implying its role in synapse formation of FS interneurons.

Previous burst kinetics studies have established some factors that may regulate burst kinetics, such as transcription factor binding^61^, feedback regulations^62^, enhancer-promoter interactions^63^, gene length, and promoter motives^19^. Luo et al.^62^ revealed that TATA-boxes can augment burst frequency in the presence of positive feedback. Larsson et al.^19^ found genes with only TATA-box or TATA-box and initiator elements had higher burst sizes in primary mouse fibroblast cells. We found that burst frequency was linked to promoter motives; TATA-box regulated genes had significantly higher burst frequencies than genes with other promoter regions in PFC neurons. These results may suggest that burst kinetics regulation is a complex mechanism and may even differ between cell types.

Summarizing our results obtained by alternative processing of FS and Pyr cell transcriptomics data, we report here about the transcriptional burst frequency and burst size parameters inferred from transcriptome-wide scRNA-seq data. We revealed that transcriptional burst kinetics is linked to a group of measures characterizing copy number distributions of the neuronal transcriptome. In addition, genes of differential burst kinetics overlapped weakly with genes of differential copy numbers. Because it is the first application of burst kinetics modeling on neuronal cell-types, we suggest the use of transcriptome-wide burst kinetics studies in efforts to discover the regulatory and potentially modulatable roles of transcriptional burst parameters in neurons.

## DATA AVAILABILITY

This paper analyzes existing, publicly available data. Single-cell RNA sequencing data analyzed in this study is found at GEO database with GSE135060 identifier. All original code has been deposited at Zenodo repository at doi:10.5281/zenodo.11092195, and is publicly available as of the date of publication. Any additional information required to reanalyze the data reported in this paper is available from the lead contact upon request.

## AUTHOR CONTRIBUTIONS

Vanda Tukacs: Investigation, Formal analysis, Visualization, Methodology, Writing—original draft. Eva Kristine Fladhus: Formal analysis, Data curation, Investigation, Methodology. Dániel Mittli: Conceptualization, Writing—review & editing. Magor László Lőrincz: Writing—review & editing. James Eberwine: Writing—review & editing. Katalin Adrienna Kékesi: Supervision, Project administration, Writing—review & editing. Gábor Juhász: Conceptualization, Supervision, Writing—review & editing.

## ACKNOWLEDGEMENTS

We thank Mária Ercsey-Ravasz for performing the hierarchical clustering of pyramidal cells and fast-spiking interneurons in this study.

## FUNDING

This study was supported by the National Research, Development and Innovation Office of Hungary Grants: [grant numbers 2017-1.2.1-NKP-2017-00002, FIEK_16-1-2016-0005] to [VT, DM, GJ, and KAK]; and National Brain Research Program NAP 3.0 of the Hungarian Academy of Sciences: [grant number NAP2022-I-3/2022] to [VT, DM, JG, and KAK], and [grant number NAP2022-I-7/2022] to [MLL].

## CONFLICT OF INTEREST

The authors declare no competing interests.

## Notes

https://github.com/vanix-21/Differential_transcription_kinetics_of_PFC_cell_types/tree/main

### Competing Interest Statement

The authors have declared no competing interest.

