## Supplemental figures for "Neuronal phenotype defined by transcriptome-wide bursting kinetics in pyramidal and fast-spiking cells"

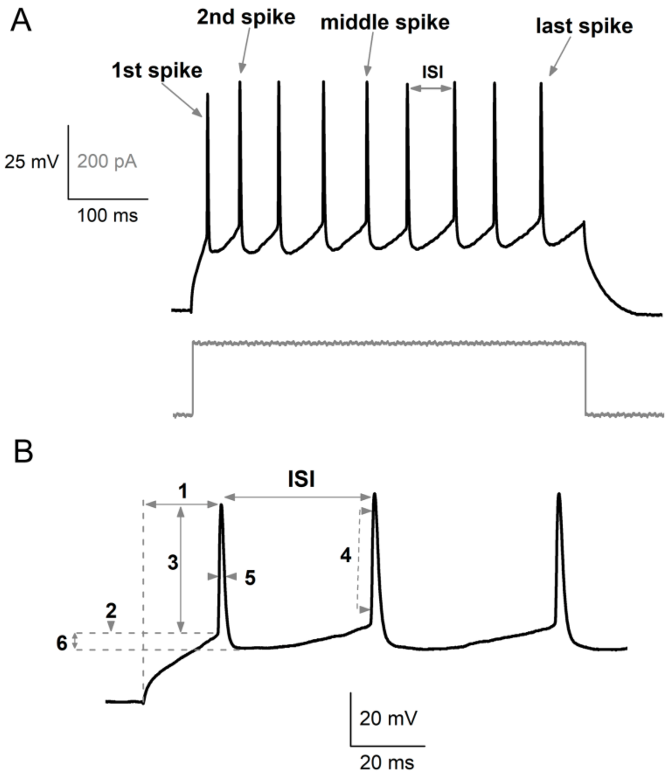


Supplementary figure S1. Parameters for cell clustering by electrophysiological data related to Figure 1. (A) The electrophysiological response of a pyramidal cell evoked by a depolarizing current step. The figure highlights the first, second, middle, and last spikes, as well as the interspike intervals (ISI). (B) The following parameters were analyzed for the first, second, and middle spikes. 1: time, 2: threshold, 3: amplitude, 4: 10-90 rise, 5: half width, 6: after hyperpolarization (AHP). The firing frequency was calculated based on the ISI between the first and second spikes and the average ISI between the second and last spikes.

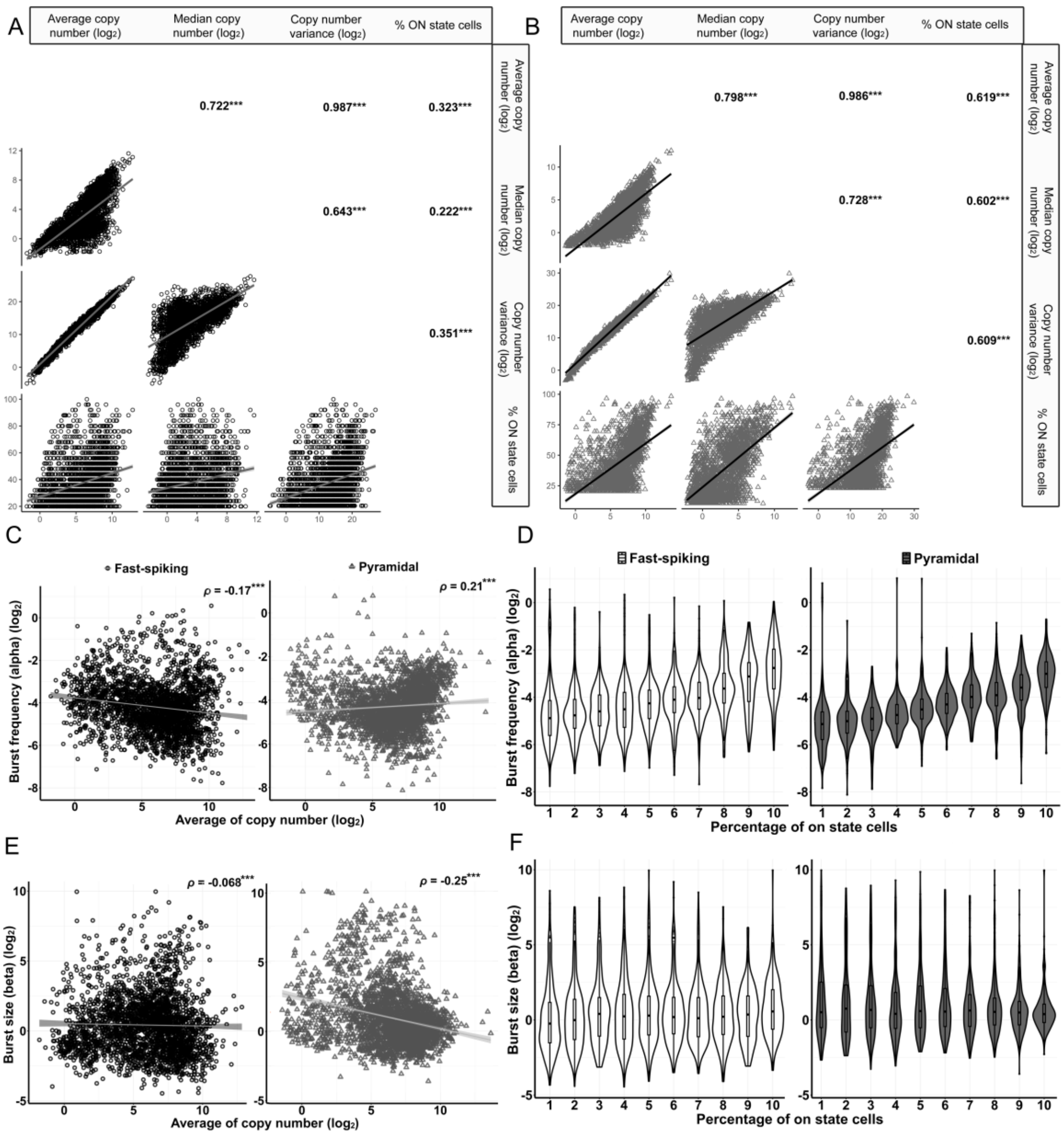


Supplementary figure S2 Correlations of copy number descriptive statistical measures and inferred burst kinetic parameters related to Figure 3. Copy number descriptive statistical measures show (A) low to high correlations in FS cells and (B) moderate to high correlations in Pyr cells.(C, D) Burst frequency and (E, F) burst size, inferred by beta-Poisson model, do not correlate with the averaged copy number and the percentage of ON state cells in FS and Pyr cells. Correlation of average copy number with (C) burst frequency and (E) burst size is low in both cell types based on Spearman's correlation coefficient. A trend of increased (D) burst frequency in relation to increasing percentage of ON state cells is shown in FS and Pyr cells; however, this trend is not apparent in the case of (F) burst size (on state frequency was divided into 10 equal bins; boxes correspond to 1st and 3rd quartile, median is indicated by line; whiskers extend to minimum and maximal values but no longer than 1.5*inter-quartile range). (Spearman's rank correlation coefficients ($\rho$) are shown on plots; ***: $p$ - value < 1.0e-03)

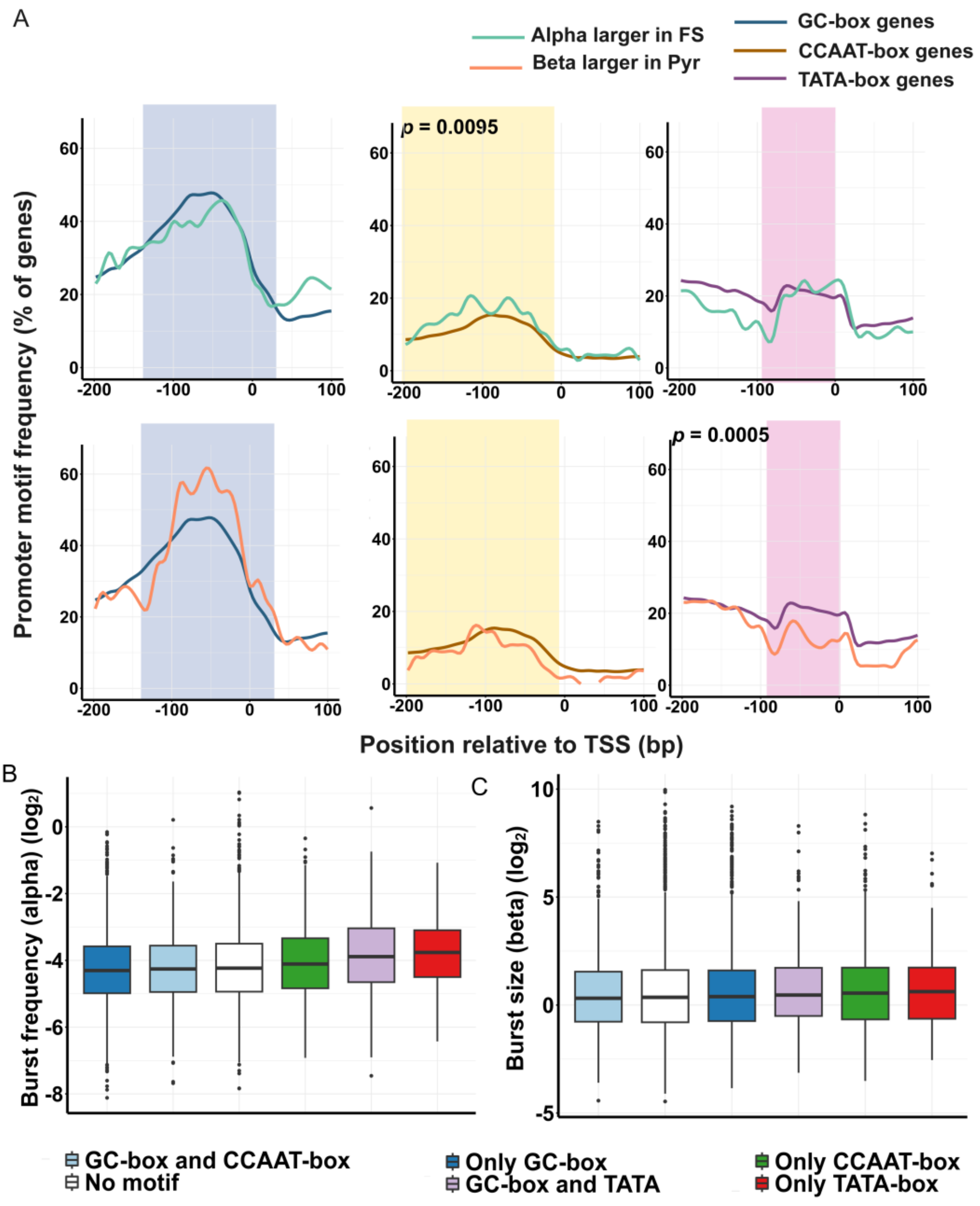


Supplementary figure S3. Promoter motif dependence of transcriptional burst kinetics related to Figure 6. (A) Relevant regions of promoter frequency distribution of genes were compared between gene groups of differential burst kinetics (Kolmogorov-Smirnoff test; significant *p*-values are shown on plot; TSS: transcription start site). Distributions of genes with only one or a combination of two promoter motives showed significant differences by Kruska-Wallis test in the case of (B) burst frequency and (C) burst size. (Boxes correspond to 1st and 3rd quartile, median is indicated by line; whiskers extend to minimum and maximal values but no longer than 1.5*inter-quartile range.) (CCAAT-box and TATA-box combination was excluded due to low number of genes in this group)
